# Structure-Based Design of Small-Molecule Inhibitors of Human Interleukin-6

**DOI:** 10.1101/2024.12.26.630392

**Authors:** Ankit Joshi, Zhousheng Xiao, Shreya Suman, Connor Cooper, Khanh Ha, Rupesh Agarwal, James A. Carson, L. Darryl Quarles, Jeremy C. Smith, Madhulika Gupta

## Abstract

Human Interleukin-6 (hIL-6) is a pro inflammatory cytokine that binds to its receptor, IL-6Rα followed by binding to gp130 and subsequent dimerization to form a hexamer signaling complex. A critical inflammation mediator, hIL-6 is associated with a diverse range of diseases and monoclonal antibodies are in clinical use that either target IL-6Rα or hIL-6 to inhibit signaling. Here, we perform high throughput structure-based computational screening using ensemble docking for small molecule antagonists for which the target conformations were taken from 600 ns long molecular dynamics simulations of the apo protein. Prior knowledge of the contact sites from binary complex studies and experimental work was incorporated into the docking studies. The top 20 scored ligands from the *in silico* studies after post analysis were subjected to *in vitro* functional assays. Among these compounds, the ligand with second-highest calculated binding affinity showed experimentally ∼84% inhibitory effect on IL6-induced STAT3 reporter activity at 10^-5^ molar concentration. This finding may pave the way for designing small molecule inhibitors of hIL-6 of therapeutic significance.

## 1. Introduction

Human interleukin-6 (hIL-6) is a multifunctional cytokine, the release of which is triggered by leukocytes, adipocytes, fibroblasts, endothelial cells, keratinocytes, and other cytokines such as IL-1 and tumor necrosis factor (TNF) as an immunologic response to infections and tissue injuries. hIL-6 transmits signals within cells by binding to either its specific receptor, IL-6Rα, or a natural soluble form of this receptor, sIL-6Rα to form a binary complex that binds to the ubiquitously expressed glycoprotein coreceptor gp130, also referred to as IL-6R [1,2]. Signal transduction occurs when the ternary complex IL-6/IL-6Rα/gp130 dimerizes to form a hexamer comprising two molecules each of IL-6, IL-6Rα, and gp130. This signaling complex activates Janus kinases and tyrosine kinases (JAK1, JAK2, TYK2), as well as signal transducer and activators of transcription (STAT), in both the JAK/STAT and RAS/MAPK pathways [3,4].

The binding of hIL-6 to IL-6Rα is referred to as ‘classical’ signaling and is responsible for the role of hIL-6 in host defence mechanisms and the stimulation of hepatocytes to produce acute-phase proteins [5]. In contrast, the binding of hIL-6 to sIL6-Rα is referred to as ‘trans’ signaling and is accountable for the immediate inflammatory reactions that occur in chronic conditions such as rheumatoid arthritis, Crohn’s disease, Castleman disease, and inflammatory bowel disease [5]. Trans signaling also mediates tumor microenvironments leading to malignancies such as multiple myeloma, colon, and pancreatic cancers [6,7]. The IL-6/ sIL-6R complex can disrupt bone equilibrium by stimulating the formation of osteoclasts resulting in abnormal bone loss, heightened fragility, and the development of arthritic conditions [8]. IL-6 in combination with TNF-α facilitates the calcification of vascular cells to induce persistent low-grade systemic inflammation in chronic kidney disease (CKD) and progression of exacerbated diabetic nephropathy (DN) [9]. Patients with different neoplastic disorders have been found to have elevated concentrations of IL-6, associated with a higher risk of cardiovascular mortality [9,10]. Recent research has also highlighted the significance of hIL-6 in severe instances of COVID-19, where it is a vital mediator of the cytokine storm syndrome that can lead to adult respiratory distress syndrome (ARDS) [11]. Thus, hIL-6 has been implicated in the pathophysiology of multiple diseases and this makes it an important therapeutic target.

Investigative approaches for therapeutic development so far have involved either reducing the formation of hIL-6 or inhibiting the binding of hIL-6 to IL-6Rα by targeting hIL-6 or the receptor. One of the popular antibodies on the market, Tocilizumab, targets IL-6Rα to treat rheumatoid arthritis, juvenile idiopathic arthritis, and Castleman disease [12]. This drug demonstrates superior effectiveness compared to methotrexate in the treatment of juvenile idiopathic arthritis and reduces hepcidin levels [13,14]. Tocilizumab demonstrated efficacy in patients with rheumatoid arthritis who did not respond to TNF antagonist therapy [15]. However, attempts to repurpose Tocilizumab for multiple myeloma and kidney transplantation showed inconsistency in results and raised safety concerns [16]. Sarilumab is an approved drug for rheumatoid arthritis and Castleman disease that targets IL-6Rα [17], while Vobarilizumab is under phase 3 clinical trials for rheumatoid arthritis [12]. Siltuximab, an approved hIL-6 antibody to treat multicentric Castleman disease has been observed to decrease iron-related complications in patients [18,19]. Further, Ancrile et al. [20] showed that the growth of Ras-driven tumors could be suppressed by targeting hIL-6 using specific antibodies. Olamkicept selectively targets trans signaling for inflammatory bowel disease and is under phase 2 trials [21]. Another inhibitor of the complex is sgp130Fc protein that is formed by fusing two sgp130 molecules to human IgG1-Fc [21–23] based on the fact that sgp130 protein functions as an intriguing natural antagonist of the hIL-6/sIL-6Rα complex [24]. However, excess of sgp130 may adversely interfere with the classical hIL-6 signaling [25,26]. Thus, the pleiotropy of hIL-6 makes it particularly challenging to develop specific inhibitors that selectively target the pro-inflammatory pathway of hIL-6.

Small molecule JAK1 inhibitors such as tofacitinib and baricitinib have been approved for rheumatoid arthritis but lack specificity and pose an increased risk of viral respiratory tract infection [27]. Other small molecule JAK1 inhibitors such as fligotinib and upadacitinib are in phase 3 clinical trials for rheumatoid arthritis and inflammatory bowel disease [28–31]. LMT-28 is a novel synthetic IL-6 inhibitor that suppresses phosphorylation of STAT3, gp130, and JAK2 and reduces IL-6-dependent TF-1 cell proliferation in mice models [32].

Despite the involvement of hIL-6 in plethora of diseases, the available drugs on the market that directly interact with the protein are antibodies limited to treating rheumatoid arthritis and Castleman disease. It may be speculated that the role of hIL-6 in these two conditions is well-defined as increased concentration levels of hIL-6 are directly linked to the progression of the disease [15,33].

It is evident that the general challenges involved in the development of small molecule inhibitors for hIL6 that target specific diseases are multifold. Different diseases involving hIL6 have differences in the pathology and the associated symptoms that need to be studied and characterized clearly. The situation is further complicated by the variations in phenotype of the affected individuals who may show varied responses for the same drug with side effects ranging from mild to severe [34,35]. Since the biologics in clinical use are confined to monoclonal antibodies that target hIL-6 or IL-6Rα, the development of economically viable small molecule inhibitors with the added advantage of ease of administration can be a boon to treating diseases involving hIL-6. Clearly, there is an unmet need to develop orally administered small molecule hIL-6 inhibitors, and we postulate that this may be achievable using ‘rational’ structure-based drug design. The research efforts directed in the developmental strategies of such potential drug lead molecules can be an asset to target the multiple diseases involving hIL-6 and improve upon existing drugs, with the possibility of enhanced efficacy and safety profiles.

Experimental high throughput screening is an expensive method for finding active small-molecules that also suffers from hit rates well below 0.1% [36]. A more effective and economical approach is to use structure-based computations in combination with assays of selected molecules. A rational approach to developing small compounds to target hIL-6 leverages understanding of the interactions that drive the formation of the key complex, hIL-6/IL-6Rα in the hIL-6 signaling cascade. The binding of hIL-6 to IL-6Rα is of paramount importance in the signaling pathway of hIL-6 as gp130 molecules cannot directly bind to hIL-6 in the absence of IL-6Rα. Thus, contact sites on hIL-6, as elucidated from hIL-6/IL-6Rα binding studies, can be targeted to identify potential lead compounds. Our previous study using extensive molecular dynamics simulations revealed that the binding of hIL-6 to IL-6Rα reduces the rigidity of residues 48 to 58 in the flexible AB loop region due to disruption of hydrogen bonds [37]. In contrast, residues 59-78 of the AB loop region lose their plasticity and become more rigid by forming contacts with the receptor on binding. Thus, the binding of hIL-6 to IL-6Rα induces structural and dynamic changes in the AB loop region of hIL-6 that subsequently facilitate the formation of hIL-6/IL-6Rα for further binding to gp130. In addition, the residues comprising helix D in hIL-6 are involved in stabilizing the interactions with the IL-6Rα primarily by two salt bridge interactions. Thus, the residues present in both the AB loop region and helix D facilitate the formation of the hIL-6/IL-6Rα complex via an interplay of electrostatic, hydrophobic, hydrogen bonding, and aromatic stacking interactions.

Previous studies have used pharmacophore modelling, docking, and structure-activity relationships to identify suitable hits for hIL-6 [38–40]. However, a comprehensive analysis of the changes induced in hIL-6 on binding with the receptor using molecular dynamics simulations and selective targeting of specific binding sites to design small molecule antagonists has not been done yet. Moreover, the docking studies till date on hIL-6 have been performed using the crystal structure and the docking algorithms may not sample all the different conformational states that the protein can have which play a crucial role in antigen-antibody recognition involving hIL-6 [38–41]. In this work, we use the conformations sampled from the molecular dynamics simulations of apo hIL-6 of Ref. 37 to perform ensemble docking on various database sets to identify potential antagonists of hIL-6. After performing several drug likeness checks and scrutinizing for PAINS, the top twenty scored compounds from *in-silico* studies were subjected to an *in vitro* functional assay. The ligand with the second highest binding affinity from docking calculations was observed to show ∼84% inhibition of hIL-6 and thus represents a lead compound for further development.

### 2. Materials and methods

hIL-6 has 186 amino acid residues and a distinct secondary and tertiary structure shown in Figure 1(a). The protein consists of four α-helices: A (Ser23-Cys46), B (Glu82-Phe107), C (Glu111 to Ala132), and D (Gln158-Met186) with three interhelical loops, AB (Glu41-Asn81), BC (residues 108-110), and CD (Lys133-Ala155). The CD loop is made up of loop CE (Lys133-Asp142) and a short mini-helix E (Asp142-Ala155). The helices A and B are oriented in the same direction while helices C and D are opposite. The two pairs of disulfide bonds formed by Cys46 (helix A)-Cys52 (AB loop) and Cys75 (AB loop)-Cys85 (helix B) stabilize the interhelix connection in hIL-6. The sites involved in binding of hIL-6 to IL-6Rα and gp130 are shown in Figure 1(b). As proteins are highly flexible molecules, it is useful to take their flexibility into account while performing docking studies. Thus, instead of docking to only a crystal structure, we consider different conformations of the protein and the ligand, in ensemble docking [42].

**Figure 1(a).**
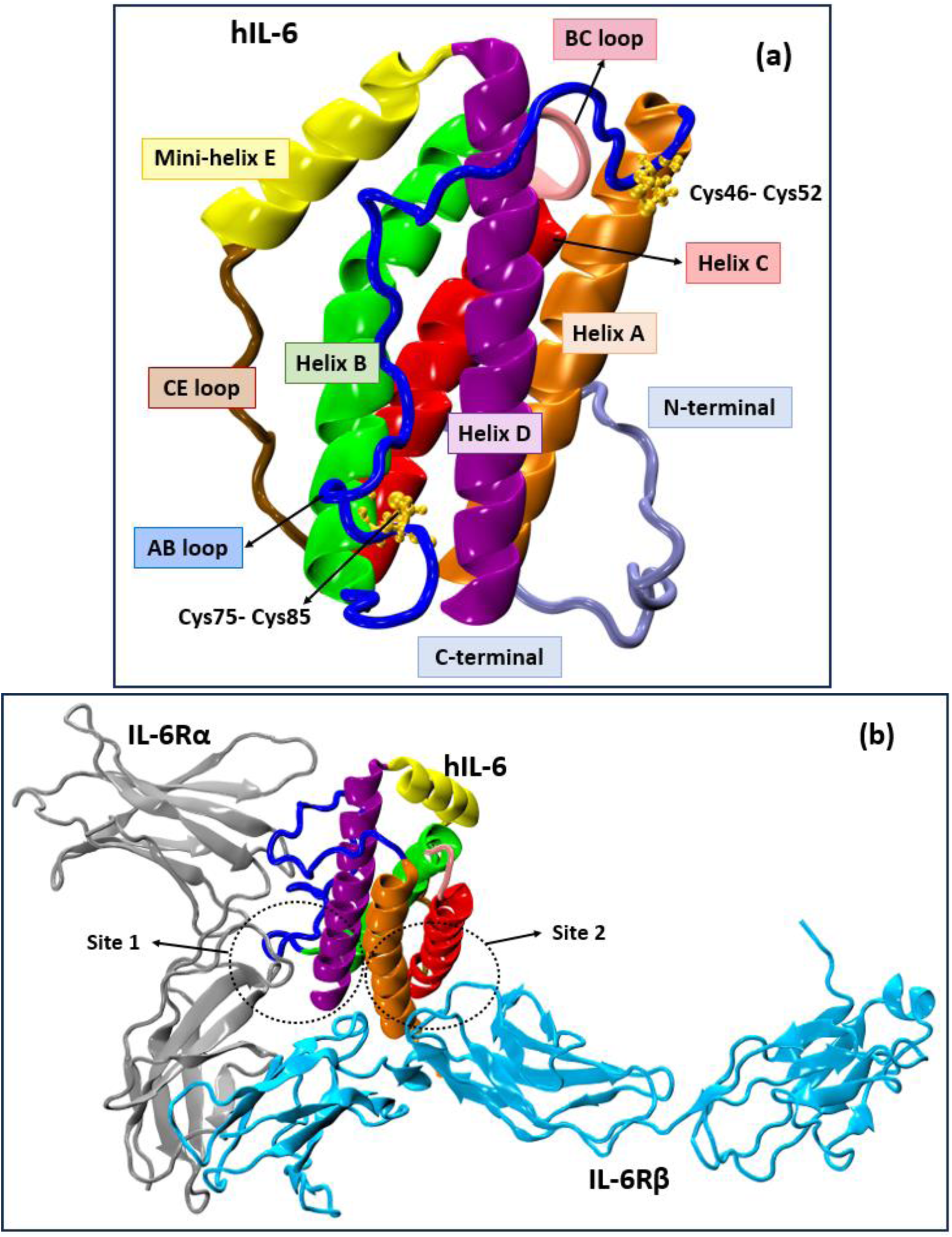
Structure of hIL-6, (b) Crystal structure of hIL-6/IL-6Rα/gp130 complex (PDB id:1P9M). The helices A, B, C, and D of hIL-6 are shown in orange, green, red, and purple, respectively. CD loop comprises mini-helix E (yellow) and CE loop (brown), while blue and pink depict loops AB and BC of hIL-6.

## 2.1 Homology Modelling

PDB IDs 1ALU (apo hIL-6) and 1P9M (hIL-6/hIL-6Rα/gp130 complex)[43,44] were used to perform homology modelling using ROSETTA [45] to simulate the absent residues in the crystal structure of hIL-6. Both, 1ALU and 1P9M lacked clear electron density for the 18 N-terminal residues. 1ALU has higher resolution than 1P9M but had 8 residues missing in the AB loop region compared to 3 residues for ALU. Consequently, a weight of 1 and 0 was assigned to 1ALU and 1P9M, respectively, during the construction of the structures. The highest-scoring model was utilized to initiate molecular dynamics simulations.

## 2.2 Molecular Dynamics (MD) Simulations

The hIL-6 modeled structure was solvated with TIP3P water [46] and neutralized with one K^+^ using CHARMM-GUI [47]. All-atom MD simulations were carried out using a GPU-accelerated version of NAMD 2.13 with the CHARMM36 force field [48,49].

The conjugate gradient method was used to perform minimization of the system followed by heating up to 300 K in the constant temperature and pressure ensemble (NPT). The equilibration and production run with total run length of 600 ns was performed in the canonical NVT ensemble. The Langevin dynamics method [50], with a damping coefficient of 1ps^-1^ was used to maintain the temperature. A Nose-Hoover barostat [50] with an oscillation period of 1ps^-1^ was used to maintain the pressure at 1 atm. A cut-off of 12 Å was used for van-der Waals, with a switch distance of 10 Å. The electrostatic interactions were computed using the Particle Mesh Ewald (PME) [51] algorithm, with a grid spacing of 1 Å. The SHAKE algorithm [52] was used to restrict the lengths of all bonds involving hydrogen atoms. All simulations were performed using an integration time step of 2 fs. The configurations in the production runs were dumped at intervals of 2 ps. Three sets of independent molecular dynamics simulations, each for 600 ns, were performed for apo hIL-6 (cumulative 1800 ns). The reported values of root mean square deviations and root mean square fluctuations showed less than 1% error over the three simulations using the standard error formula [37]. Thus, the run length of 600 ns was adequate to sample the conformational space of hIL-6, and one of the simulations was used in the next section to generate the different ensembles for docking.

## 2.3 Ensemble docking algorithm to select distinct conformations of hIL-6 for screening

Ensemble docking studies were performed here using MD simulations of apo hIL-6 for 600 ns [37]. The KFC2 server [53,54] was used to calculate hotspots for the hIL6/IL-6Rα complex to identify key contact sites for binding in the crystal structure (1P9M) as well as in our simulations. The results reveal that Phe76, Gln77, Gln177, Leu180, Arg181, Ala182 and Arg184 of hIL-6 were the predicted hotspots. These data are in excellent agreement with the experimental studies suggesting the role of residues 178-186 of hIL-6 in protein-receptor binding [55–58]. This information was incorporated into our clustering protocol to obtain the conformationally different structures. We used root mean square deviations (RMSD) of residues within 5 Å of residues 178-186 to cluster MD-derived protein structures using the GROMOS method in GROMACS [59] with a cutoff of 2 Å. Using RMSD = 2.5 Å resulted in only 3 different conformations of hIL-6 that may be too less to represent different conformational states of hIL-6. It may be noted here that the cut-off value for clustering should be chosen to obtain an optimal number of different conformations of the protein. A high cut-off value for RMSD would give few conformations leading to inadequate representation of different conformations possible for the protein. A low cutoff value would result in too many conformations of the protein that may not be too different from each other to yield different binding energies in docking. Thus, a cutoff of 2 Å was chosen in this work to have a good representation of the different conformations for hIL-6 to account for its flexibility while docking.

## 2.4 Docking protocol and selection of database for docking

The program VinaMPI, a high throughput docking version of AutodockVina, was used to speed up the calculations [60] and targeted docking was performed using prior information about the binding site in hIL-6 [37,55,61]. The NCI database (2,65,242 compounds) [62], Enamine diversity set (∼50,000 compounds) [63], Enamine PPI library (40,640 compounds) [64], and SWEETLEAD database (4,030 approved drugs, herbal isolates, and medicines) [65] were used to perform high throughput screening to identify potential ligands for targeting hIL-6. OPEN BABEL and MGLTools were used to generate structure files [66,67]. A box of 30 Å × 30 Å × 30 Å centered around the 178-186 residues of hIL-6 was chosen for docking. The exhaustiveness parameter in AutodockVina was set to 50.

## 2.5 Selection criteria for choosing compounds for in vitro screening

The ligands were arranged in the order of their docking scores and were subjected to drug likeness checks. A PAINS filter was also used to screen out compounds with sub structural features that make them promiscuous compounds in high throughput screening. Lipinski’s rule of five, which was introduced in 1997, is commonly referred to as a guideline for identifying drug-like compounds. Its main purpose is to tackle the issue of low solubility and permeability in oral medications. This is achieved by ensuring that the molecular weight (MW) is less than 500, the lipophilicity (cLogP) is less than 5, the number of hydrogen bond donors (OH + NH groups) is less than 5, and the number of hydrogen bond acceptors (O + N atoms) is less than 10 [68–70]. The Lipinski rule of five was applied to assess the drug-likeness of the top-scoring ligands [71]. A strict implementation of the Lipinski’s rule has been observed to pose a barrier in some cases to exploring novel drugs [72]. Therefore, here if fewer than two rules are violated, a non-zero drug-likeness value was assigned to the ligand. The ligands were also checked for the absence of any reactive segments based on such as metals, N/O/S-N/O/S single bonds, thiols, acyl halides, Michael acceptors, azides, esters and others [73]. Compounds that formed only hydrophobic interactions or had less than 2 hydrogen bonds were removed from the list. The top 20 compounds obtained thereafter were subjected to *in vitro* experimental studies.

## 2.6 In Vitro Functional Assays

The 20 top scored compounds were purchased from Enamine. HEK293T cells were cultured in Dulbecco’s modified Eagle’s medium (DMEM) containing 10% fetal bovine serum and 1% penicillin and streptomycin (P/S). For IL6-mediated activation of the IL-6Rα/gp130/STAT3 singling, HEK293T cells were transiently transfected with either empty expression vector or full-length human IL-6, along with IL-6Rα, gp130, and STAT3 (Addgene, Inc.) luciferase reporter system and Renilla luciferase-null as internal control plasmid. Transfections were performed by electroporation using Cell Line Nucleofector Kit R according to the manufacturer’s protocol (Amaxa Inc.). Thirty-six hours after transfection, the cells were treated with the test compound at 10^−5^ M or in a range of 10^−8^-10^−4^ M and vehicle only was served as the control. After 5 hours, the cells were lysed, and luciferase activities were measured using a Synergy H4 Hybrid Multi-Mode Microplate Reader and Promega Dual-Luciferase Reporter Assay System (Madison, WI). Three independent experiments were run for each scenario to obtain sufficient statistics. Statistical significance between two groups was evaluated by unpaired the 2-tailed t-test and that between multiple groups was evaluated by one-way analysis of variance (ANOVA) with the Turkey multiple comparison test. These calculations were

### 3 Results and discussion

*In-silico studies*: High-throughput virtual ensemble docking was performed on the conformations derived from extensive MD simulations with the goal of identifying candidate small molecules binding to hIL-6 that prevent the formation of the hIL-6/IL-6Rα complex. The chemical structures of the remaining top 20 ligands with the highest binding affinities as calculated with Autodock Vina are shown in Figure 2. All the 20 ligands are highly hydrophobic and mostly aromatic. Table 1 lists physical properties related to drug likeness of the top docked compounds. The molecular weights of all the reported compounds are less than 500. All the 20 compounds have logP < 5 except Z445038774. The other rules relating to the number of hydrogen bond donors and acceptors were also followed for the top docked ligands. Interestingly, all the compounds listed in Table 1 are found to be from the Enamine database. This emphasizes the importance of small, well-curated targeted libraries in binding proteins in certain specific cases. Moreover, since hIL-6 interacts with the receptor at the interface of the helix rather than having a binding pocket or cavity, it may be speculated that the enamine diversity set can be used for inhibiting protein-protein interactions compared to NCI and SWEETLEAD databases. Table 1 also shows topological polar surface area, tPSA, which is the key parameter to predict drug permeability and binding affinity. Compounds with high tPSA values (>140 Å²) are typically less likely to be orally bioavailable.

**Figure 2.**
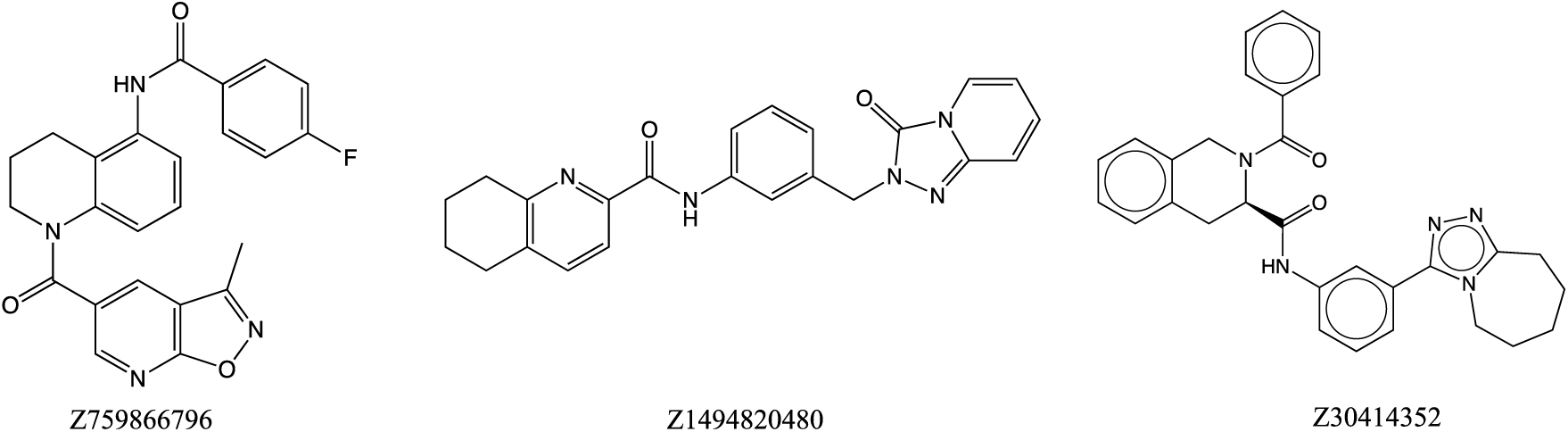
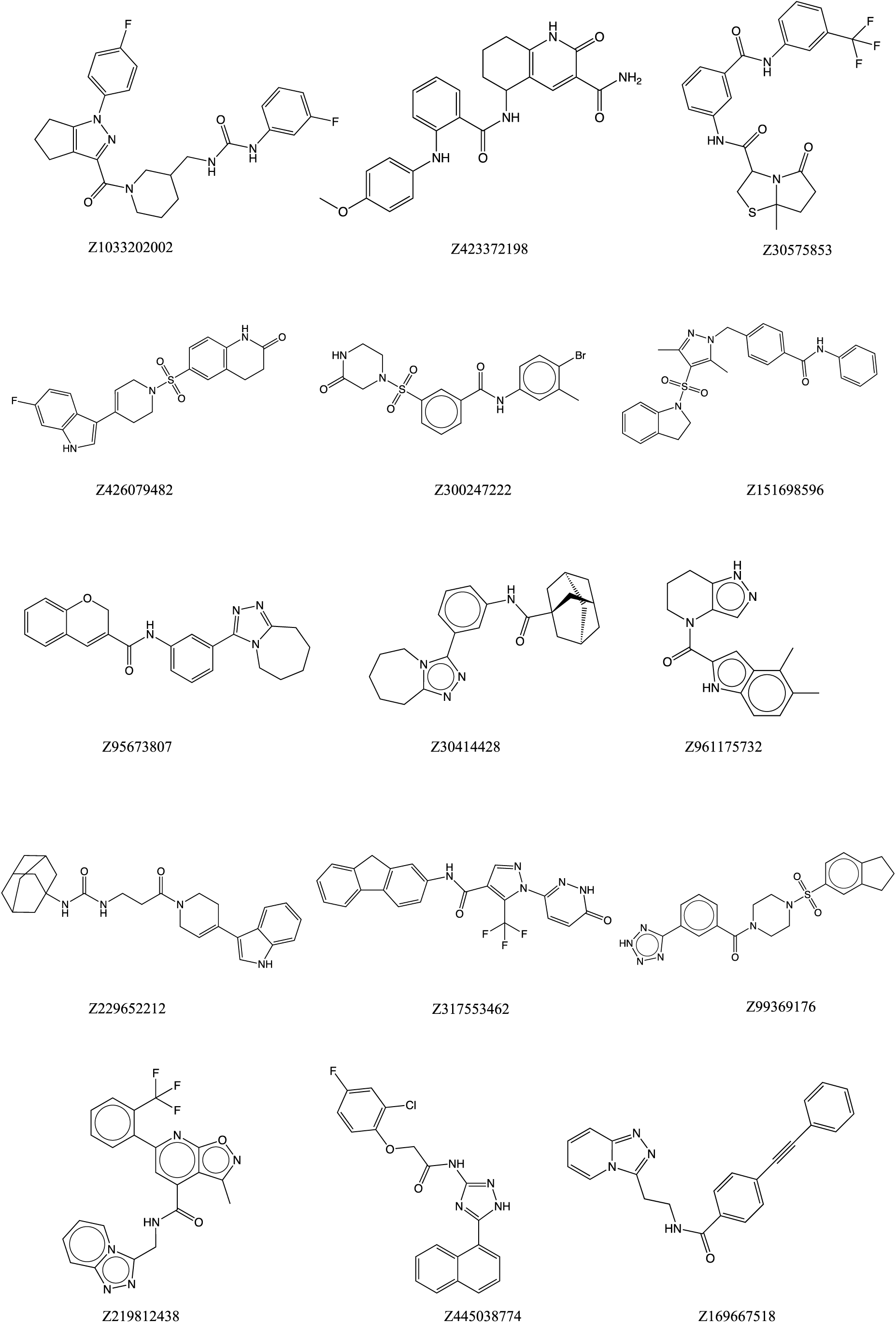
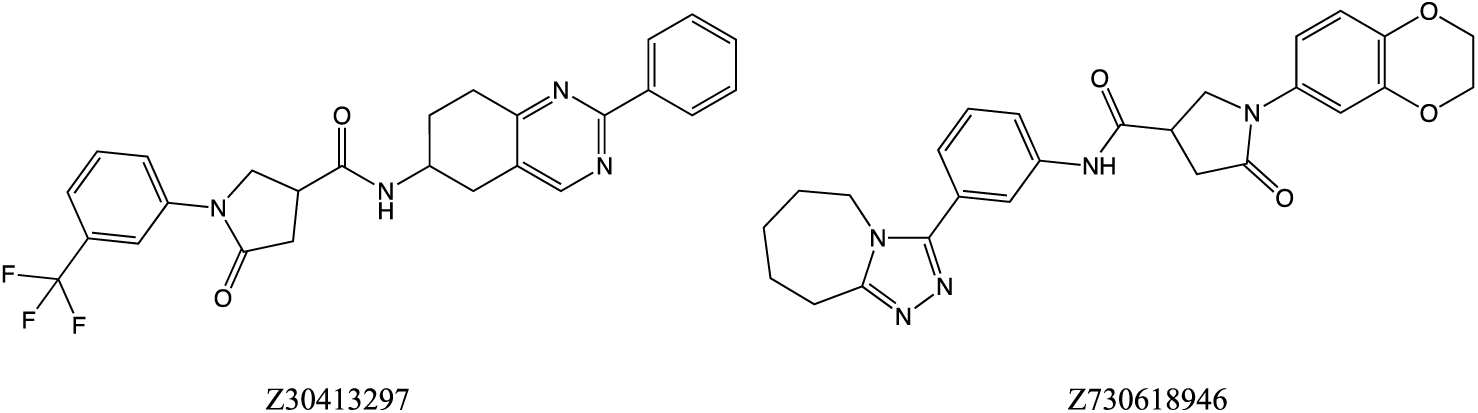
Chemical structures of top 20 scored ligands from Autodock Vina. performed using GraphPad Prism 5.0 (San Diego, CA). The IC50 of Z169667518 was obtained graphically from concentration-effect curves using GraphPad Prism 5.0 (San Diego, CA).

**Table 1.** Drug likeness properties [75] of top 20 ligands with the highest docking scores computed from AutoDock Vina^a^.

| Compound ID | Mol. Formula | Mol. Wt. | logP | Net charge | H-bond donors | H-bond acceptors | tPS A | Docking Score (kcal/mol) |
| --- | --- | --- | --- | --- | --- | --- | --- | --- |
| Z229652212 | C<sub>27</sub>H<sub>34</sub>N<sub>4</sub>O<sub>2</sub> | 446.595 | 4.442 | 0 | 3 | 2 | 77 | -9.150 |
| Z169667518 | C<sub>23</sub>H<sub>18</sub>N<sub>4</sub>O | 366.424 | 3.102 | 0 | 1 | 4 | 59 | -8.881 |
| Z30414428 | C<sub>24</sub>H<sub>30</sub>N<sub>4</sub>O | 390.531 | 4.826 | 0 | 1 | 4 | 59 | -8.840 |
| Z423372198 | C<sub>24</sub>H<sub>24</sub>N<sub>4</sub>O<sub>4</sub> | 432.48 | 3.033 | 0 | 4 | 5 | 126 | -8.774 |
| Z30575853 | C<sub>22</sub>H<sub>20</sub>F<sub>3</sub>N<sub>3</sub>O<sub>3</sub>S | 463.481 | 4.35 | 0 | 2 | 4 | 78 | -8.711 |
| Z219812438 | C<sub>22</sub>H<sub>15</sub>F<sub>3</sub>N<sub>6</sub>O<sub>2</sub> | 452.396 | 4.19 | 0 | 1 | 7 | 98 | -8.701 |
| Z95673807 | C<sub>23</sub>H<sub>22</sub>N<sub>4</sub>O<sub>2</sub> | 386.455 | 4.086 | 0 | 1 | 5 | 69 | -8.627 |
| Z1494820480 | C<sub>23</sub>H<sub>21</sub>N<sub>5</sub>O<sub>2</sub> | 399.454 | 3.07 | 0 | 1 | 6 | 81 | -8.618 |
| Z30414352 | C<sub>30</sub>H<sub>29</sub>N<sub>5</sub>O<sub>2</sub> | 491.595 | 4.877 | 0 | 1 | 5 | 80 | -8.611 |
| Z730618946 | C<sub>26</sub>H<sub>23</sub>F<sub>3</sub>N<sub>4</sub>O<sub>2</sub> | 480.49 | 4.189 | 0 | 1 | 4 | 75 | -8.582 |
| Z30413297 | C<sub>26</sub>H<sub>27</sub>N<sub>5</sub>O<sub>4</sub> | 473.533 | 3.434 | 0 | 1 | 7 | 98 | -8.574 |
| Z99369176 | C<sub>21</sub>H<sub>22</sub>N<sub>6</sub>O<sub>3</sub>S | 438.513 | 1.502 | - | 1 | 6 | - | -8.287 |
| Z151698596 | C<sub>27</sub>H<sub>26</sub>N<sub>4</sub>O<sub>3</sub>S | 486.597 | 4.552 | 0 | 1 | 5 | 84 | -8.281 |
| Z759866796 | C<sub>24</sub>H<sub>19</sub>FN<sub>4</sub>O<sub>3</sub> | 430.439 | 4.516 | 0 | 1 | 5 | 88 | -8.258 |
| Z317553462 | C<sub>22</sub>H<sub>14</sub>F<sub>3</sub>N<sub>5</sub>O<sub>2</sub> | 437.381 | 4.21 | 0 | 2 | 5 | 92 | -8.238 |
| Z426079482 | C<sub>22</sub>H<sub>20</sub>FN<sub>3</sub>O<sub>3</sub>S | 425.485 | 3.67 | 0 | 2 | 3 | 82 | -8.208 |
| Z1033202002 | C<sub>26</sub>H<sub>27</sub>F<sub>2</sub>N<sub>5</sub>O<sub>2</sub> | 479.531 | 4.313 | 0 | 2 | 4 | 79 | -7.879 |
| Z961175732 | C<sub>17</sub>H<sub>18</sub>N<sub>4</sub>O | 294.358 | 3.101 | 0 | 2 | 2 | 64 | -7.772 |
| Z300247222 | C<sub>18</sub>H<sub>18</sub>BrN<sub>3</sub>O<sub>4</sub>S | 452.33 | 2.13 | 0 | 2 | 4 | 95 | -7.331 |
| Z445038774 | C<sub>20</sub>H<sub>14</sub>C<sub>1</sub>FN<sub>4</sub>O<sub>2</sub> | 396.809 | 5.084 | 0 | 2 | 4 | - | -7.280 |

<sup>a</sup>The above table lists the docking ID, molecular formula (Mol. Formula), molecular weight (Mol. Wt.), lipophilicity (logP), net charge, number of hydrogen bond donors, number of hydrogen bond acceptors, topological polar surface area(tPSA) as obtained from Zn database [75] and docking score for each ligand as obtained from AutoDock Vina.

Figure 3a depicts the calculated binding pose for the ligand with the highest calculated binding affinity in our docking studies - Z229652212. This ligand interacts with helix D and the AB loop region that together comprise site I for binding to hIL-6Rα as shown in Figures 3(b-c). The alkyl groups present in Leu66 and Leu167 interact with the ligand. Met69, Ser171, and Glu174 are involved in forming hydrogen bonds with the ligand. Pro67, Lys68, and Phe175 form carbon-hydrogen bonds with the ligand. This type of non-conventional hydrogen bonding has gained importance recently over a decade and has been observed to play a crucial role in stabilizing the protein ligand interactions [74].

**Figure 3.**
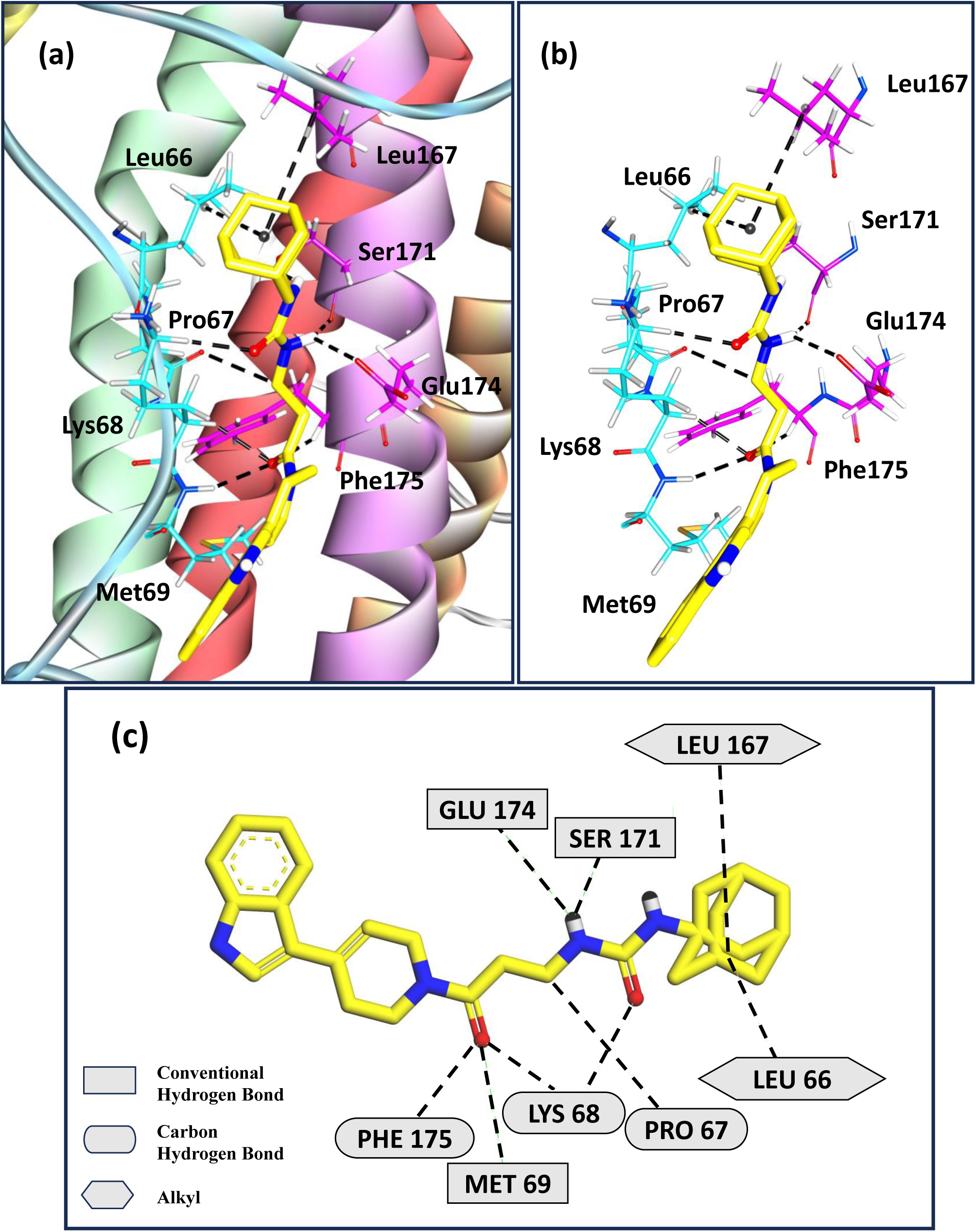
(a) Docked pose for the highest binding score ligand Z229652212 with hIL-6, (b) and (c) represent the sites of interaction of ligand Z229652212 and hIL-6.

Figure 4a depicts the binding pose for the ligand Z169667518 which shows the second highest calculated binding affinity with hIL-6. Figures 4(b,c) shows that Leu66, Pro67, Met69, and Phe76 from AB loop region, with Leu167, Ser171, and Phe175 of helix D of hIL6, interact with the ligand. Leu66, Met69, and Leu167 stabilize the ligand through alkyl interactions. Pro67, Ser171, and Phe175 form carbon-hydrogen bonds with the ligand. Phe76 further stabilizes the protein-ligand complex by 𝛑-𝛑 stacking interactions. The same residues of hIL-6 involved in stabilizing the interactions with the ligands Z229652212 and Z169667518 in this study were observed to form contacts with the receptor leading to their rigidification in our previous study [37]. In particular, Phe residues and aromatic rings play a crucial role in stabilizing the hIL6/ hIL-6Rα complex. Thus, the results from this work are consistent with our previous study [37] on the role played by the AB loop region in facilitating antigen-antibody recognition in addition to the binding site present on helix D.

**Figure 4.**
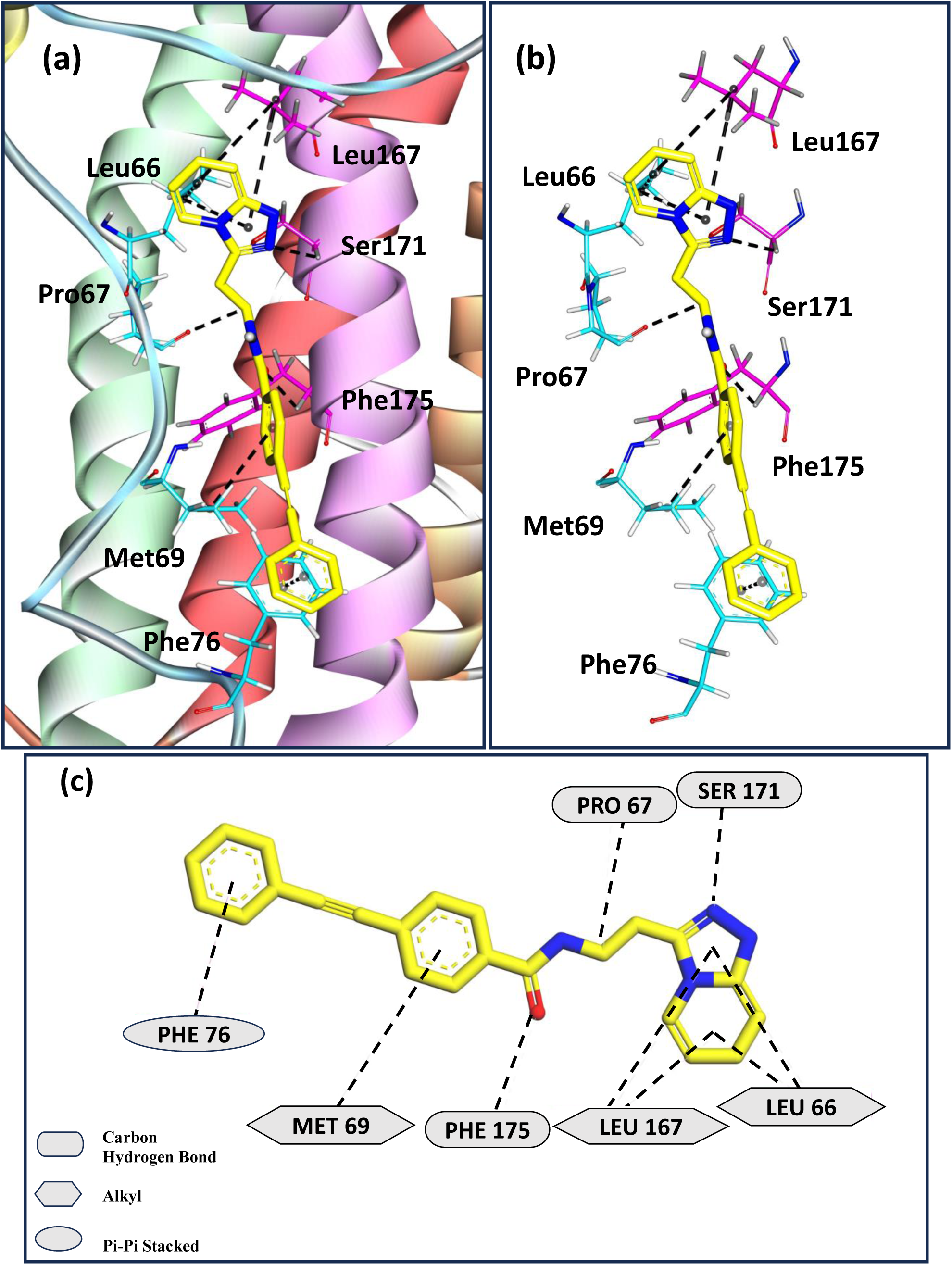
(a) Docked pose for the second highest binding score ligand Z169667518 with hIL-6, (b) and (c) represent the sites of interaction of ligand Z169667518 and hIL-6.

#### In vitro screening of top 20 calculated ligands for antagonization of IL6-mediated activation of IL6Rα/GP130/STAT3 signaling in vitro

Using the *in vitro* screening assay described in Section 2.4, we experimentally tested 20 high-scoring chemical probes identified from the *in silico* virtual screen at an initial concentration of 10 μM in the presence of IL6 as shown in Figure 5. Sixteen of the 20 compounds exhibited statistically significant measurable effects in inhibiting IL6-stimulated STAT3 reporter activity (Figure 5A). Compounds Z445038774, Z30413297, Z961175732, and Z99369176 at 10 μM concentration had no significant effect on IL6-induced signal transduction.

**Figure 5.**
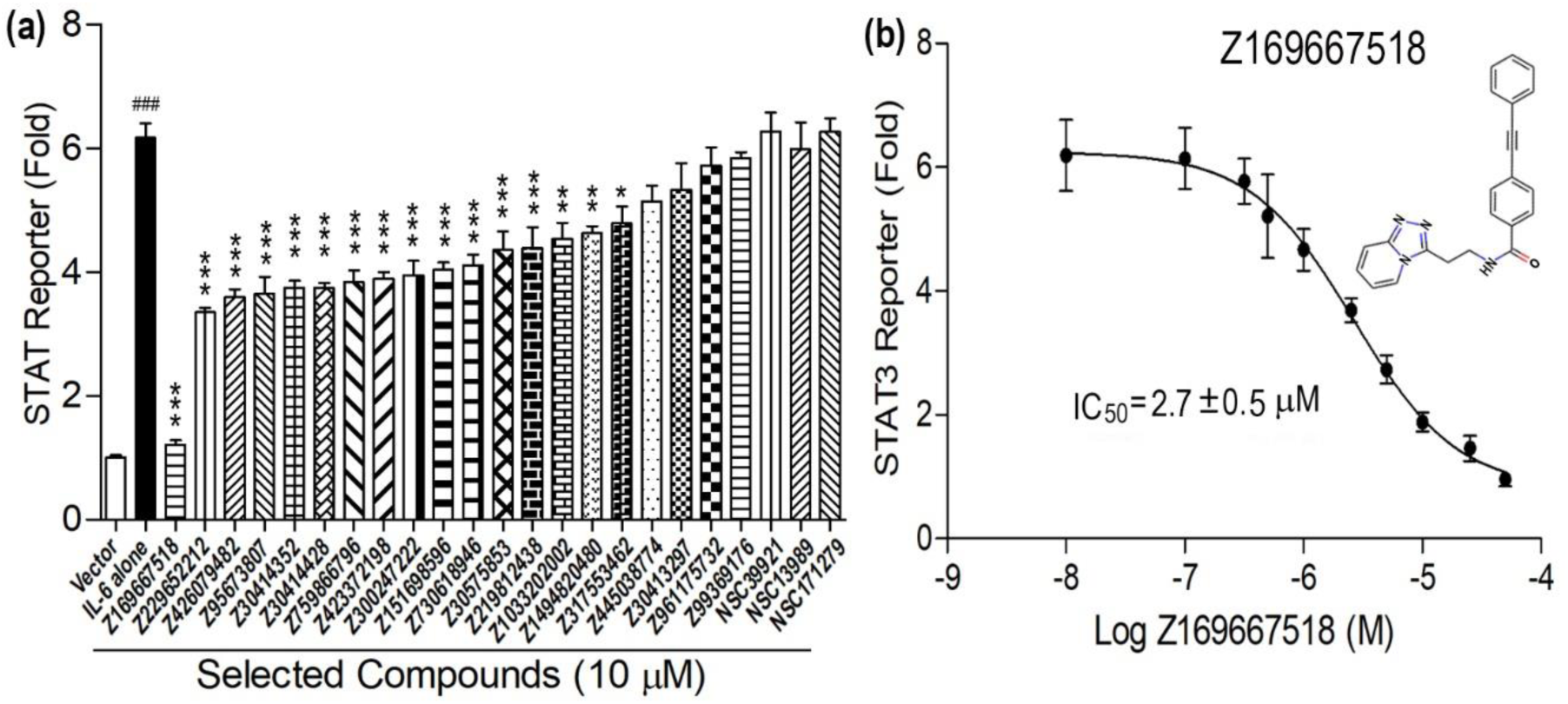
*In vitro* functional assays of the top 20 compounds from Enamine and 3 from NCIdatabase selected from the *in silico* virtual screen. (a) Effects of these compounds on IL6 induced STAT3 reporter activities in the transiently IL6Rα/GP130-transfected HEK 293T cells. (b) Dose−response curve of Z169667518 on IL6-induced STAT3 reporter activities. Data are means ± SD from three independent experiments. ^###^ indicates statistically significant difference from empty vector group. ** and *** indicate statistically significant difference from IL-6 vehicle group. *P* values were determined by one-way ANOVA with Tukey’s multiple-comparisons test.

At 10 μM concentration, 15 compounds exhibited partial (less than 50%) inhibition of IL6-induced STAT3 reporter activity while one compound, Z169667518, stood out by exhibiting ∼84% inhibitory effect (Figure 5a). In the ensemble docking studies, this compound was calculated to interact strongly with hIL6, as shown in Table 1 and Figure 4, and had the second-highest calculated binding affinity.

To explore the dose-response effects of Z169667518 we performed additional studies using doses ranging from 10^−8^ to 10^−4^ M. Z169667518 exhibited dose-dependent inhibition of IL6-induced STAT3 reporter activity (Figure 5b). The estimated median inhibitory concentration (IC50) value for Z169667518 was 2.7 ± 0.5 µM (Figure 5b). Optimization of this lead compound could potentially result in IL6/IL6Rα/GP130 interaction inhibitors with sub-micromolar to nanomolar binding affinities for IL-6.

It may be noted that despite the elimination of NCI compounds during the filtering criteria discussed in Section 2.5, three of the top-scored NCI compounds (Figure 6, Table 2) were subjected to *in vitro* screening. Figure 5 depicts that none of these compounds showed any significant effect on IL6-induced signal transduction validating their exclusion during the selection criteria as well.

**Figure 6.**
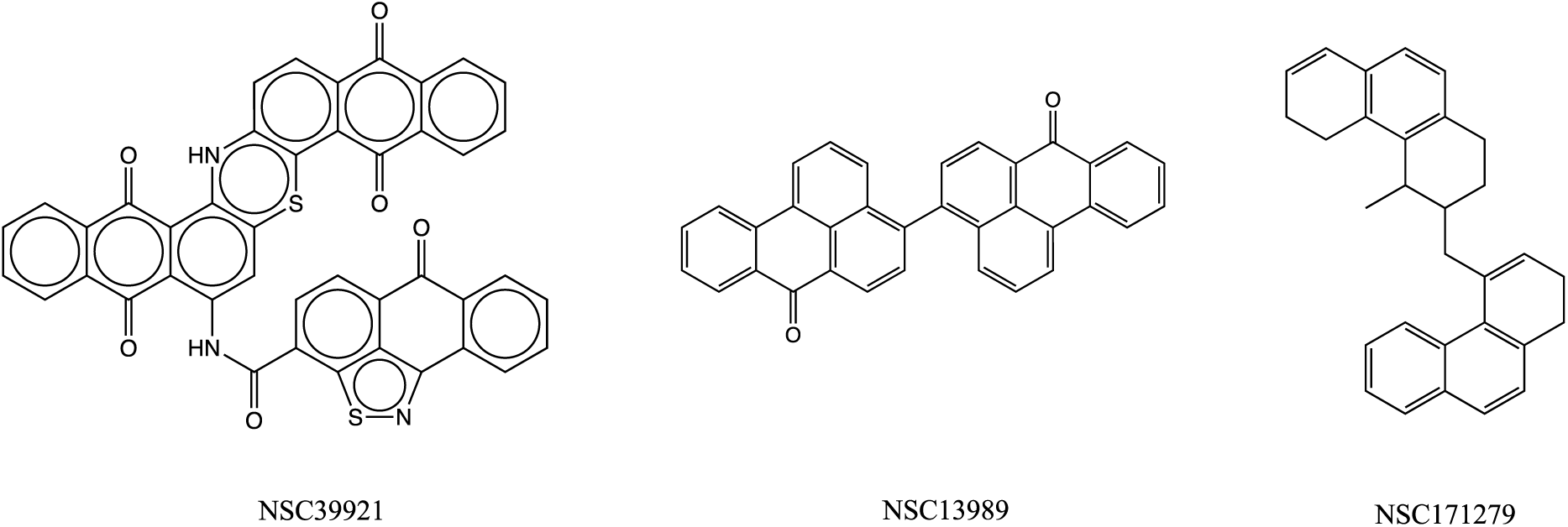
Chemical structures of top 3 scoring NCI compounds subjected to *in vitro* studies.

**Table 2.** Drug likeness properties [75] of 3 NCI ligands with the highest docking scores subjected to *in vitro* studies.

| Compound ID | Mol. Formula | Mol. Wt. | logP | Net charge | H-bond donors | H-bond acceptors | tPSA | Docking Score (kcal/mol) |
| --- | --- | --- | --- | --- | --- | --- | --- | --- |
| NSC39921 | C<sub>43</sub>H<sub>43</sub>N<sub>3</sub>O<sub>6</sub>S<sub>2</sub> | 762.0 | 6.3 | 0 | 2 | 8 | 192.9 | -11.1 |
| NSC13989 | C<sub>34</sub>H<sub>18</sub>O<sub>2</sub> | 458.5 | 8.8 | 0 | 0 | 2 | 34.1 | -10.6 |
| NSC171279 | C<sub>30</sub>H<sub>30</sub> | 390.6 | 7.1 | 0 | 0 | 0 | 0 | -10.6 |

### 4 Conclusions

Interleukin-6 is a pleiotropic, proinflammatory cytokine produced by a variety of cell types, including lymphocytes, monocytes, and fibroblasts. hIL-6 is prominent in the etiology of several diseases and the cytokine storm. Anti-IL-6 strategies [76,77] may be broadly applicable to a wide range of medical conditions such as pericarditis, gout, type 2 diabetes, and others that are still under investigation. However, hIL-6 also plays an important role in immune responses and tissue regeneration. Therefore, hIL-6 has a narrow therapeutic window and long-term inhibition can result in numerous adverse effects, such as increased vulnerability to bacterial and fungal infections. To overcome these limitations, it will be important to develop a variety of drugs that address specific functions of hIL-6.

Although therapeutic antibodies against hIL-6 have been developed, small-molecule therapeutics, which have the clear advantage of oral availability, are lacking. Here, a structure-based target site prediction approach was used, informed by understanding the detailed mechanism of formation of the IL-6/IL-6Rα complex that is pertinent in triggering the signaling cascade of hIL-6. Structural information on contact sites of hIL-6 involved in the formation of the binary complex was employed in virtual high throughput screening through targeted ensemble docking studies. The top 20 scoring ligands from enamine as shown in table 1 with 3 more NCI compound as shown in table 2 from the docking studies were subjected to *in vitro* analysis. The ligand with second highest binding affinity as shown in table 1 from *in silico* studies showed ∼84% inhibitory effect on hIL6 binding for the *in vitro* experimental assay and an IC50 value of 2.7 ± 0.5 µM.

Cell assays are excellent for identifying compounds that produce a desired phenotype, but do not confirm the proposed mechanism of action or provide details of specific target binding. For the perturbation of protein-protein interactions obtaining this type of information is non-trivial, but would be of use in lead optimization and target validation studies. Notwithstanding, the discovery of this ligand demonstrates that biophysical structure-based computational methods can be combined with assays to find small molecules capable of preventing the formation of the hIL-6/IL-6Rα complex so as to inhibit hIL6-induced STAT3 reporter activities. As such the compound found here may form the basis of lead optimization and pre-clinical work, and further studies using the present method targeting the interface would likely lead to the discovery of additional chemical probes.

### Authors contributions

A.J., S.S., and M.G. performed all the simulations, docking studies and analysis and wrote the main manuscript along with preparation of figures and tables. Z.X. and J.C. performed the *in vitro* functional assay of top 20 scored compounds. C.C. screened out the compounds for Lipinski’s rule and PAINS. K.H. was involved in running docking of few compounds. R.A. performed homology modelling of N-terminus of hIL-6 and performed filtering of certain compounds. D.Q. and J.S. provided important inputs to improve the manuscript and reviewed the manuscript.

### Data Availability Statement

The data will be made available upon request to the authors.

## Acknowledgements

MG acknowledges support from SERB-POWER grant received from Science and Engineering Research Board (SERB), under the Department of Science and Technology (DST) Government of India (Reference No. SPG/2021/002224). MG acknowledges support from Faculty Research Scheme (Reference No. FRS(160)/2021-2022/CHEMISTRY) by IIT(ISM) Dhanbad. This research used resources of the Compute and Data Environment for Science (CADES) at the Oak Ridge National Laboratory, which is supported by the Office of Science of the U.S. Department of Energy under Contract No. DE-AC05-00OR22725. This research was also supported by the United States National Institutes of Health (NIH) grants R01-DK121132 and R01-AR071930 to L. Darryl Quarles and Zhousheng Xiao.

